# Regulation of TCR-induced Transcriptional Kinetics by Interleukin 2-inducible T Cell Kinase (ITK) in CD8^+^ T cells

**DOI:** 10.1101/2025.09.22.677398

**Authors:** Josh Hunkins, Zoe K. Bedrosian, Uddeep Chaudhury, Alayna Rosales, Mary J. Michaels, Masahiro Ono, Abhyudai Singh, Leslie J. Berg

**Affiliations:** Department of Immunology/Microbiology, University of Colorado CU Anschutz, Denver, CO; Department of Life Sciences, Imperial College London, London, United Kingdom; Department of Electrical and Computer Engineering, Department of Biomedical Engineering, University of Delaware, Newark, DE

## Abstract

The initial activation of naïve CD8^+^ T cells induces three major transcription factor pathways, NFAT, NFκB, and AP-1, predominantly regulated by T-cell receptor (TCR) signaling. Downstream of the TCR, the Tec family tyrosine kinase ITK modulates the transcriptional response in T cells by differentially affecting the kinetics and magnitude of activation of these three transcription factors. How signaling through ITK regulates the duration and/or termination of TCR signaling has not been investigated. To address this, we utilized the “*Nr4a3*-Tocky” reporter mouse which provides information on the kinetics of TCR signaling over time courses from hours to days. OT-I CD8^+^ *Nr4a3-Tocky* T cells were stimulated with peptide ligands of varying affinities, and cells were assayed for a panel of surface and intracellular markers of activation and differentiation at multiple time points post-stimulation. As previously reported, at early time points ITK signaling enhanced the kinetics and magnitude of expression of these markers. At later time points the absence of ITK led to persistent expression of several proteins, indicating an important role for ITK in the termination of TCR-dependent transcriptional responses. Using a dual ITK/ RLK inhibitor (PRN694) at 24h post OVA peptide stimulation it was found that this function of ITK was required in the first 24 hours. These findings reveal a possible key negative regulatory mechanism that is programmed in the first day of activation but impacts CD8^+^ T cell gene expression patterns days later.

**One Sentence Summary Statement:** The Tec family tyrosine kinase ITK differentially regulates early and late T-cell receptor transcriptional responses.

## Introduction

The initial signals a naïve CD8^+^ T cell receives through its TCR, coupled with integrated signals from costimulatory receptors and cytokines/chemokines, regulate the differential expression of genes responsible for determining T cell fate and function (1, 2). Stimulation of the TCR leads to activation of downstream kinases that promotes transcriptional responses dominated by three transcription factor families, NFAT, NFκB, and AP-1 (3–6). An essential step in this pathway is the activation of phospholipase C-γ1 (PLCγ1), which cleaves the membrane phosphoinositide PIP_2_ into two second messengers, diacylglycerol (DAG) and IP_3_. PLCγ1 is activated by recruitment to the LAT-SLP76 adapter complex, a signaling complex rapidly assembled following TCR stimulation, and by PLCγ1 tyrosine phosphorylation (7–10). The Tec family tyrosine kinase ITK, while not essential for all PLCγ1 activation, is responsible for phosphorylating PLCγ1 and enhancing its activity, leading to increased production of DAG and IP_3_ following TCR stimulation (7, 11). DAG activates the Ras-Erk MAP-kinase pathway, ultimately leading to the induction of active AP-1. DAG also binds to and activates Protein kinase C-θ (PKCθ) to initiate the canonical NF-κB transcriptional response. The second messenger IP_3_ promotes calcium influx into the T cell, leading to calcineurin activation and the activation of NFAT (12).

Stimulation of T cells with antigens of varying affinities or at varying doses is known to impact the kinetics of T cell activation. This results from variations in the duration of TCR:pMHC binding (13–15) as well as the localization and density of TCR microclusters (16–18). Costimulatory signals, such as those mediated by CD28 or by cytokines, can also alter T cell activation kinetics (19–22). The activities of TCR proximal kinases such as Lck, ZAP-70, ITK, and the rate of LAT phosphorylation (23, 24) also contribute to the rate of T cell activation (25, 26). As a result, increases in antigen affinity have been shown to decrease the time to the initial round of T cell proliferation *in vivo* (27) and *in vitro* (28).

In our earlier studies, we examined how variations in TCR signaling translated into changes in the three transcriptional pathways downstream of PLCγ1 (1). We measured the kinetics and magnitude of NFAT and NF-kB(p65) nuclear translocation, and of ERK-MAP-kinase (Erk-MAPK) phosphorylation using OT-I TCR transgenic CD8^+^ T cells. OT-I cells recognize a peptide of chicken ovalbumin (OVA-N4 (257–264); SIINFEKL) bound to MHC class I (K^b^) with high affinity. Variants of this peptide with single amino acid substitutions at the 4th position are recognized with lower affinities by the OT-I TCR, and when used to stimulate T cells produce weaker TCR signals than those generated by recognition of the high affinity N4 peptide (27). Overall, we found that weaker TCR stimulation using the altered peptide variants SIITFEKL (OVA-T4) and SIIGFEKL (OVA-G4) caused a profound reduction in the rate of onset of downstream TCR responses (1). In addition, and consistent with previous studies (5, 25, 29), we demonstrated that both NFAT and Erk-MAPK activation show all-or-nothing ‘digital’ responses to stimulation, such that all responding cells had the same amount of nuclear NFAT or p-Erk-MAPK regardless of the dose or affinity of the stimulating antigen. In contrast, NF-kB(p65) activation was ‘graded’ with responding cells having varying amounts of nuclear p65 in correlation to the strength of the TCR signal they received. This prior study also examined the contribution of ITK signaling to the activation of NFAT, NF-kB(p65) and Erk-MAPK. We found that ITK had no impact on either the kinetics or magnitude of Erk-MAPK activation. Examination of NFAT activation showed that ITK signaling was only required when cells were stimulated with low affinity antigen, and primarily impacted the kinetics of NFAT nuclear localization and the percentage of T cells in the population that activated NFAT (1). In contrast, ITK had a major effect on NF-kB(p65) activation, affecting the magnitude of nuclear p65 evident in each cell and under all TCR stimulation conditions (1).

To date, the role of variations in TCR signal strength on the kinetics of T cell gene expression has been studied at early time-points and has focused on examining how proximal signaling events lead to alterations in the onset or initiation of T cell activation marker expression (1, 2, 14, 25–27, 30–32). However, linkage of the PLCγ1-dependent transcriptional pathways to changes in specific gene expression responses has not been examined in detail. Furthermore, as each of the downstream surface and intracellular markers commonly used to study T cell activation vary in their dependence on these early transcription factor pathways (33–36), perturbations in these pathways have heterogeneous effects on gene induction (1, 26, 37). A more comprehensive knowledge of these details would be invaluable in efforts to generate engineered T cells with predictable downstream functions *in vivo*.

While much is known about how proximal events downstream of the TCR lead to the expression of many T cell activation markers, significantly less is known about the mechanisms leading to the termination of early gene expression responses. Here we examined TCR signaling kinetics together with a panel of surface receptors and intracellular factors in wild-type and *Itk^-/-^* OT-I CD8+ T cells using the fluorescent reporter *Nr4a3*-Tocky (38). The *Nr4a3*-Tocky reporter line, which produces the Tocky fluorescent reporter protein downstream of the *Nr4a3* promoter, was developed to track temporal events following TCR signaling (38). The reporter protein, a variant of mCherry (39), emits blue fluorescence when first translated and has a half-life of ∼4.1 hours before undergoing a time dependent spectral conversion to emit red fluorescence. This reporter allows cells with recent TCR signaling (blue fluorescent) to be distinguished from cells with sustained signaling (blue + red fluorescent) and from cells that have terminated/arrested signaling (red fluorescent). *Nr4a3* transcription is highly correlated with CD3/CD28-mediated signaling, and is regulated by both the NFAT and Erk-MAPK pathways (38, 40–42) making it an excellent temporal metric for TCR signaling. Utilizing this system, along with small molecule inhibitors of the NFAT, NF-kB, and Erk-MAPK pathways, we examined the expression of a panel of proteins following activation of CD8+ T cells. These data demonstrated the multifactorial regulation of most early T cell activation markers, and the importance of TCR signal strength in dictating the kinetics of gene expression responses. Further, we found that ITK signaling is critical for the early dynamics of a potential negative feedback pathway that tunes the expression of many of these genes, and that this negative feedback mechanism is induced within the first 24 hours of T cell stimulation. These findings indicate that ITK regulates the kinetics of TCR-mediated gene expression at both the initiation and the termination phases of the response.

## Results

### TCR signal strength determines the duration and magnitude of signaling and the expression of T cell activation markers

To test the effects of TCR signal strength on the overall expression kinetics of CD8^+^ T cells, we stimulated OT-I *Rag*1^-/-^ *Nr4a3-*Tocky splenocytes *in vitro* with 1nM OVA peptides for 5, 12, 24, 48, and 72 hours. Two peptide ligands at two doses were used: OVA-N4 (high affinity) and OVA-T4 (lower affinity), which vary in their potency at activating OT-I CD8^+^ T cells (27). Overlaying all time points stimulated with 1nM OVA on a single plot showed increased proportions of Tocky-Blue^+^ Tocky-Red^+^ cells in the OVA-N4 stimulated sample compared to OVA-T4, along with elevated Median Fluorescent (MFI) Intensity of Tocky-Blue at 24h and 48h post-stimulation (Fig. 1A-C). In both the 1nM OVA-T4 and OVA-N4 stimulated cells, peak expression levels for CD69 were at 12h, whereas *Nr4a3*-Tocky Blue, CD44, CD25, PD-1, IRF4, and IRF8 peaked at 24h based on the normalized MFI (Fig. 1D). For cMyc and Egr2, the highest levels of expression were at 5h. Comparing cells stimulated with OVA-N4 to those stimulated with OVA-T4 revealed differences in the peak magnitudes of induced expression for most proteins, but in general, little impact on the general pattern of expression over the course of three days (72h) (Fig. 1D).

**Fig. 1.**
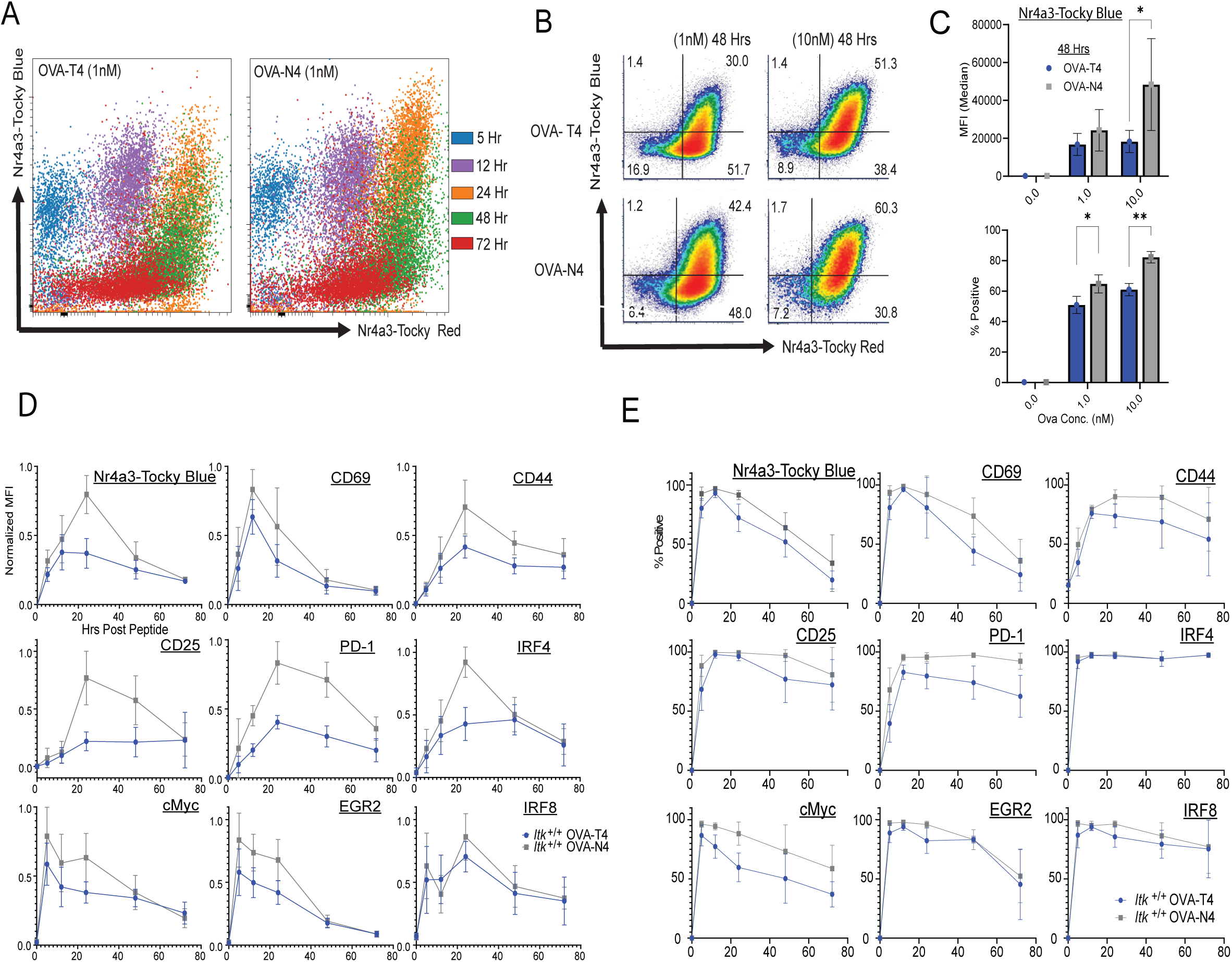
TCR stimulation with high affinity pMHC promotes prolonged signaling and prolonged expression of multiple T cell activation markers. OT-I *Nr4a3*-Tocky splenocytes were stimulated with OVA-N4 or OVA-T4 at 1nM or 10nM and examined at 5, 12, 24, 48 and 72h post-stimulation. (A) Tocky-Blue versus Tocky-Red fluorescence in cells stimulated with 1nM of each peptide are overlaid for all 5 time points. (B) Representative flow plots of Tocky-Blue versus Tocky-Red expression at 48h. (C) Compilation of data showing percentages of Tocky-Blue-positive cells and Tocky-Blue MFI at 48h. Data are means +/- SD of 6-8 biological replicates and 5 independent experiments. Multiple paired t-test *p ≤ 0.05, **p ≤ 0.01, ***p ≤ 0.001, ****p ≤ 0.0001. (D, E) Compiled timecourse data for cells stimulated with 1nM OVA-N4 (grey) or OVA-T4 (blue) showing MFIs of expression in cells positive for the indicated marker at each time (D) or percentages of marker positive cells at each timepoint (E). Means +/-SD of normalized MFI (calculated as % experimental max) or percentages of positive cells are shown from 6-8 biological replicates of 5-6 independent experiments.

However, a closer analysis of the Tocky-Blue signal, a surrogate for ongoing TCR stimulation, showed that a greater proportion of CD8⁺ T cells activated with OVA-N4 continued to express Tocky-Blue at 48h compared to cells stimulated with OVA-T4 peptide (Fig. 1B-C). Based on the known half-life of the Tocky-Blue signal, the presence of Tocky-Blue-positive cells at this time indicated that TCR signaling was persisting in those cells for a minimum of 40 hours. OVA-N4 stimulated cells showed a greater percentage of CD8 T cells remain positive for CD69, CD44, CD25, PD-1, and cMyc compared to OVA-T4 stimulated cells at 48h, reinforcing the observation that higher affinity TCR stimulation leads to a modest increase in the duration of expression of several activation markers (Fig. 1E).

### TCR signal strength tunes the dynamics of initial activation signals that broadly impact gene expression

To assess the effect of ligand affinity and abundance on the initial onset of TCR signaling, we examined the OT-I *Nr4a3*-Tocky CD8+ T cells over the first 5h of activation using Tocky-Blue versus Tocky-Red fluorescence in OVA-T4 and OVA-N4 stimulated cells, along with a panel of additional early activation markers. Both the percentage of positive cells as well as the relative expression levels, shown as MFI, among positive cells were quantified for each protein. Examination of Tocky-Blue and CD69 (an Erk-MAPK-dependent early activation marker (43)) co-expression showed that at a population level the two proteins increased in parallel following stimulation with either peptide (Fig. 2A), despite the slower kinetics of the response to OVA-T4 (Fig. 1A). Transcription factors that are also expressed such as c-Myc, EGR2, and IRF4 also showed faster kinetics of expression as peptide affinity and/or peptide dose increased (Fig. 2B). A similar finding was observed for the expression of the high affinity subunit of the IL-2 receptor, CD25 (Fig. 2C). A compilation of data for a set of surface markers and transcription factors showed the consistent delay in upregulation of these responses after 5 hours of TCR stimulation when OT-I cells were stimulated with the lower affinity OVA-T4 variant compared to OVA-N4 (Fig. 2D).

**Fig. 2.**
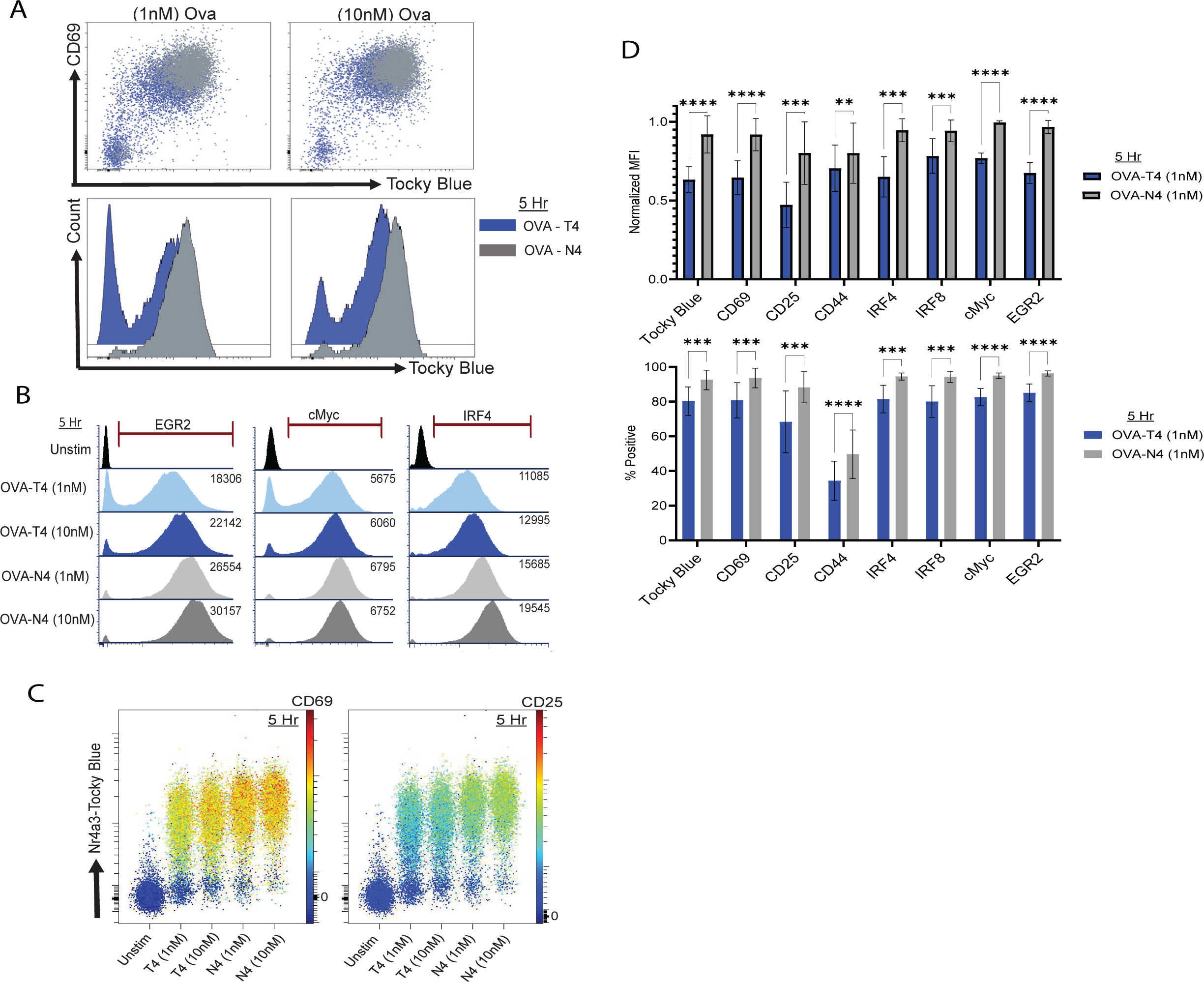
Stimulation of CD8^+^ T cells with reduced TCR signal strength delays the upregulation of multiple surface markers and transcription factors. OT-I *Nr4a3*-Tocky splenocytes were stimulated *in vitro* with OVA-N4 or OVA-T4 at 1nM or 10nM for 5h. (A) Pseudocolor plots of Tocky-Blue versus CD69 are shown with OVA-N4 stimulated cells (Grey) overlaid on OVA-T4-stimulated cells (Blue), with histograms of Tocky-Blue fluorescence below. (B) Histograms of intracellular EGR2, cMyc, and IRF4 expression; MFI of positive cells, gated as shown, on right. (C) Comparative pseudocolor plots showing expression of Tocky-Blue fluorescence on Y-axis with CD69 or CD25 fluorescence on a colored continuous scale as shown. (D) Compilation of data showing normalized MFI (% experimental max at 5h; top) and percentages of positive cells (bottom) for the indicated proteins. Data are from 6-8 biological replicates of 5 independent experiments. Statistics derived from multiple paired t-test, *p ≤ 0.05, **p ≤ 0.01, ***p ≤ 0.001, ****p ≤ 0.0001.

### *Itk^-/-^* CD8^+^ T cells show prolonged expression of several early activation markers independent of variations in TCR signal strength

To assess the duration of TCR signaling and the impact of ITK on the persistence of T cell activation, we used *Nr4a3*-Tocky Blue expression as a measure of ongoing TCR signaling (38). Along with Tocky-Blue, we measured a set of surface and intracellular proteins in *Itk*^+/+^ and *Itk^-/-^*T cells over 5, 12, 24, 48, and 72 hours following stimulation with OVA-N4 or OVA-T4 peptide. This allowed us to compare the kinetics of expression of cell surface receptors associated with activation as well as of transcription factors.

We first examined Tocky-Blue versus Tocky-Red signals along with CD69 expression levels at each time point and observed prolonged expression of CD69 beginning at 24h of stimulation in *Itk*^-/-^ compared to wild-type (*Itk^+/+^*) cells (Fig. 3A). Examining the expression of CD69 Tocky-Blue/Red single positive (SP), double positive (DP), and double negative events at 48h post-stimulation, we found that *Itk^-/-^* CD8^+^ T cells showed a greater percentage of CD69^+^ cells compared to *Itk*^+/+^ cells with both OVA-T4 and OVA-N4 stimulation (Fig. 3B). Expanding this analysis to other proteins demonstrated that *Itk^-/-^*T cells showed prolonged expression of many proteins at 48h and 72h compared to *Itk*^+/+^ T cells. Tocky-Blue, CD69, CD44, cMyc, and IRF4 expression all remained elevated in *Itk^-/-^* T cells at 48h post-stimulation compared to *Itk*^+/+^ cells (Fig. 3C); a similar pattern was observed for IRF8, Eomesodermin (Eomes), CD25, and PD-L1 (Supp. Fig 1). In the case of IRF4, protein levels increase in *Itk^-/-^* T cells between 24h and 48h, whereas the opposite was the case for *Itk*^+/+^ cells indicating there was prolonged upregulation of IRF4 in cells lacking ITK (Fig. 3C). In addition, for cells stimulated with OVA-N4, *Itk^-/-^* T cells often showed a greater percentage of cells retaining expression of induced proteins at 48h compared to *Itk*^+/+^ cells (Fig. 3C). Overall, this analysis demonstrated that the absence of ITK from the TCR signaling pathway does not mimic the effects of reducing TCR signal strength seen by lowering peptide affinity or peptide dose in Fig. 1D. Instead, the absence of ITK produces a diminished early response, but leads to prolonged expression of activation markers at later time points (>24h).

**Fig. 3.**
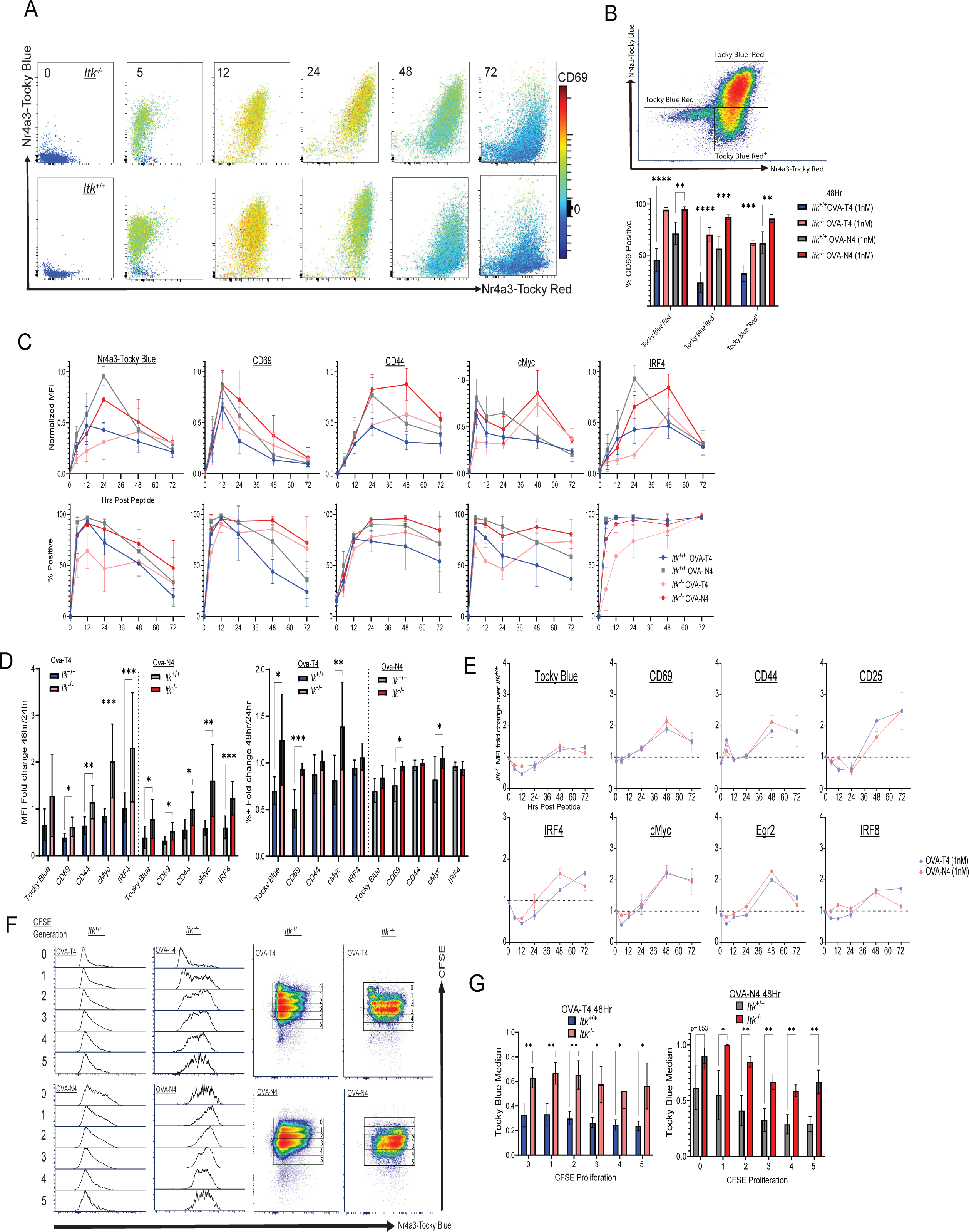
Impaired termination of activation gene expression in the absence of ITK. (A) 0-72h timecourse of Tocky-Blue versus Tocky-Red expression for *Itk^-/-^* (top) and *Itk*^+/+^ (bottom) OT-I *Nr4a3*-Tocky cells stimulated with OVA-N4 at 1nM. CD69 expression is represented as colored continuous on each density plot. (B) Compilation of data comparing percentages of CD69-positive cells at 48h on gated Tocky Blue^+^Red^+^, Tocky Blue^-^Red^+^ and Tocky Blue^-^Red^-^ populations, with gating scheme shown on representative density plot. Statistical analysis performed with 2-way ANOVA with Tukey’s correction. Bars are representative of 3-5 experiments with 3-8 biological replicates showing mean +/-SD; *p ≤ 0.05, **p ≤ 0.01, ***p ≤ 0.001, ****p ≤ 0.0001. (C) Compilation of data from 0-72h time courses show normalized MFI (% experimental max for marker; top) and percentages of positive cells (bottom). (D) The ratio of each MFI at 48h/24h is plotted; bars <1 indicate a higher signal at 24h than 48h, and bars >1 indicate a higher signal at 48h compared to 24h. Statistics derived from multiple unpaired t-test, *p ≤ 0.05, **p ≤ 0.01, ***p ≤ 0.001, ****p ≤ 0.0001. (E) The ratio of the MFI in *Itk^-/-^* cells over *Itk*^+/+^ cells is plotted over time for each marker. Points >1 indicate a higher signal in *Itk^-/-^* cells versus *Itk^-/-^*cells and points <1 indicate a higher signal in *Itk*^+/+^ cells compared to *Itk^-/-^* cells. Data are mean +/-SD from 3-8 biological replicates across 3-5 experiments. (F) *Itk*^+/+^ and *Itk^-/-^*cell were labeled with CFSE and stimulated with (1nM) OVA-T4 and (1nM) OVA-N4 for 48h. Histograms of Tocky-Blue fluorescence for cells in each division cycle are shown. (G) Normalized MFI (ratio of MFI to max experimental MFI at 48h) of Nr4a3-Tocky Blue fluorescence for cells in each division cycle at 48h, comparing *Itk^+/+^* and *Itk^-/-^* cells stimulated with Ova-T4 (1nM) and Ova-N4 (1nM). Statistics derived from unpaired t test; *p ≤ 0.05, **p ≤ 0.01, ***p ≤ 0.001, ****p ≤ 0.0001 of 3-4 biological replicates from 3 experiments.

To visualize these findings in a more systematic manner, we calculated fold change of MFI as well as percentages of positive cells for each protein at 48h versus 24h (Fig. 3D). In these graphs values of 1 indicate equivalent expression at 48h compared to 24h, whereas values <1 indicate reduced expression at 48h relative to 24h. This analysis indicates that for a set of proteins, including Tocky-Blue, CD69, CD44, cMyc and IRF4, *Itk^-/-^*OT-I cells show an increased ratio of expression per cell (48/24h) compared to *Itk*^+/+^ cells. In several cases, ratios of the percent positive cells are also higher in *Itk^-/-^* cells, most notably Tocky Blue, CD69 and cMyc in response to stimulation with low affinity OVA-T4 peptide (Fig. 3D).

To better understand the relative effect of peptide affinity on this persistence, we calculated the ratios of MFIs for multiple proteins in *Itk^-/-^* versus *Itk*^+/+^ OT-I cells over the entire 72 hour time-course (Fig. 3E). In these graphs, data points >1 indicate that the expression levels of that protein are higher in *Itk^-/-^* compared to *Itk*^+/+^ cells at a given time point, and data points <1 indicate higher expression in *Itk*^+/+^ compared to *Itk^-/-^*. At early time points, *Itk*^+/+^ cells express higher levels of several proteins compared to *Itk^-/-^* cells, indicated by data points <1. This is particularly evident for cells stimulated with OVA-T4, confirming previous findings (26, 44) that the contribution of ITK to TCR-dependent responses is substantially greater under conditions of weak TCR signaling (44). These graphs also confirm the finding that at later time points (>24h), *Itk^-/-^*cells express higher levels of these proteins than *Itk*^+/+^ T cells, indicated by data points >1. Furthermore, this phenomenon is not dependent on TCR signal strength as the ratio of activation marker expression in *Itk^-/-^* over *Itk*^+/+^ cells after stimulation with either high or low affinity antigen is similar for most proteins examined (Fig. 3E).

To rule out the possibility that our observations of prolonged TCR signaling and extended duration of activation marker expression in *Itk^-/-^*cells were due to differences in cell proliferation, we assessed *Nr4a3*-Tocky Blue expression across cell divisions using CFSE dilution. *Itk*^+/+^ and *Itk^-/-^* OT-I cells were stimulated for 48h with 1nM OVA-N4 or OVA-T4 and then analyzed for Tocky-Blue signal within each cell division cycle. This analysis confirmed that, compared to *Itk*^+/+^ cells, *Itk^-/-^* cells express persistent Tocky-Blue with both high and low TCR stimulation at each corresponding cell division cycle (Fig. 3F,G). These data indicate that persistent *Nr4a3*-Tocky Blue expression in *Itk^-/-^*cells is not due to reduced Tocky Blue dilution resulting from lower proliferation of *Itk^-/-^* cells (45, 46).

Finally, to validate our findings of prolonged protein expression in CD8+ T cells lacking ITK and link *Nr4a3* expression to promoter activity post peptide stimulation, we developed a simple mathematical model based on a one-dimensional ordinary differential equation. This model leverages the known ∼4.1 hour half-life for the conversion of Tocky Blue into the red fluorescent form. Based on this approach, the inferred *Nr4a3* promoter activity and the corresponding model-predicted dynamics of the Tocky-Blue signal are shown (Supp. Fig 2). With 1nM OVA-N4 stimulation the promoter activity in *Itk*^+/+^ cells peaks at 12h post stimulation. The same peptide stimulation in *Itk^-/-^* cells also led to a similar timing of peak promoter activity but the peak was reduced by 30% compared to *Itk*^+/+^. Notably, the ensuing decrease in promoter activity is much slower in *Itk^-/-^*as compared to *Itk*^+/+^ (Supp. Fig 2). Stimulation with OVA-T4 exhibits a striking difference in promoter activity between the two different genotypes. In *Itk*^+/+^ cells, promoter activity peaks at 4h followed by a decline. In contrast, in *Itk^-/-^* cells the promoter activity plateaus after a rapid initial increase, and remains relatively increased until 60h post stimulation, after which it exhibits a sharp decline. This sustained promoter activation in *Itk^-/-^* cells explains the rising Tocky-Blue signal even up to 50h post 1nM T4 stimulation and validates the prolonged phenotype we see at the protein expression level.

### Inhibition of ITK after 24 hours of stimulation does not alter the kinetics or duration of T cell activation responses

We next sought to determine the timing of the requirement for ITK in terminating the responses to T-cell stimulation. To assess this, we stimulated *Itk*^+/+^ OT-I CD8^+^ T cells with low affinity OVA-T4 for 24h, washed off the peptide and cultured the cells for an additional 24h in the presence or absence of the small molecule dual inhibitor of ITK/RLK, PRN694 (47). Cells were analyzed by flow cytometry at 24h, prior to PRN694 addition, and at 48h. As shown, addition of PRN694 at 24h post-stimulation did not lead to prolonged expression of CD69 or CD44 (Fig. 4A). As a control, we compared *Itk^+/+^*OT-I cells cultured for the entire 48h with PRN694 and *Itk^-/-^*OT-I cells stimulated for 48h in the absence of PRN694, and verified that the presence of PRN694 during the first 48h of stimulation produced similar prolonged expression observed with *Itk^-/-^* OT-I cells when stimulated with OVA-T4 (Fig 4B). These data demonstrated that the effects of ITK on gene expression kinetics resulted from ITK activity in the first 24h post-stimulation, as prolonged expression of multiple T cell activation markers was not observed when the ITK inhibitor was added to cells after the first 24h.

**Fig. 4.**
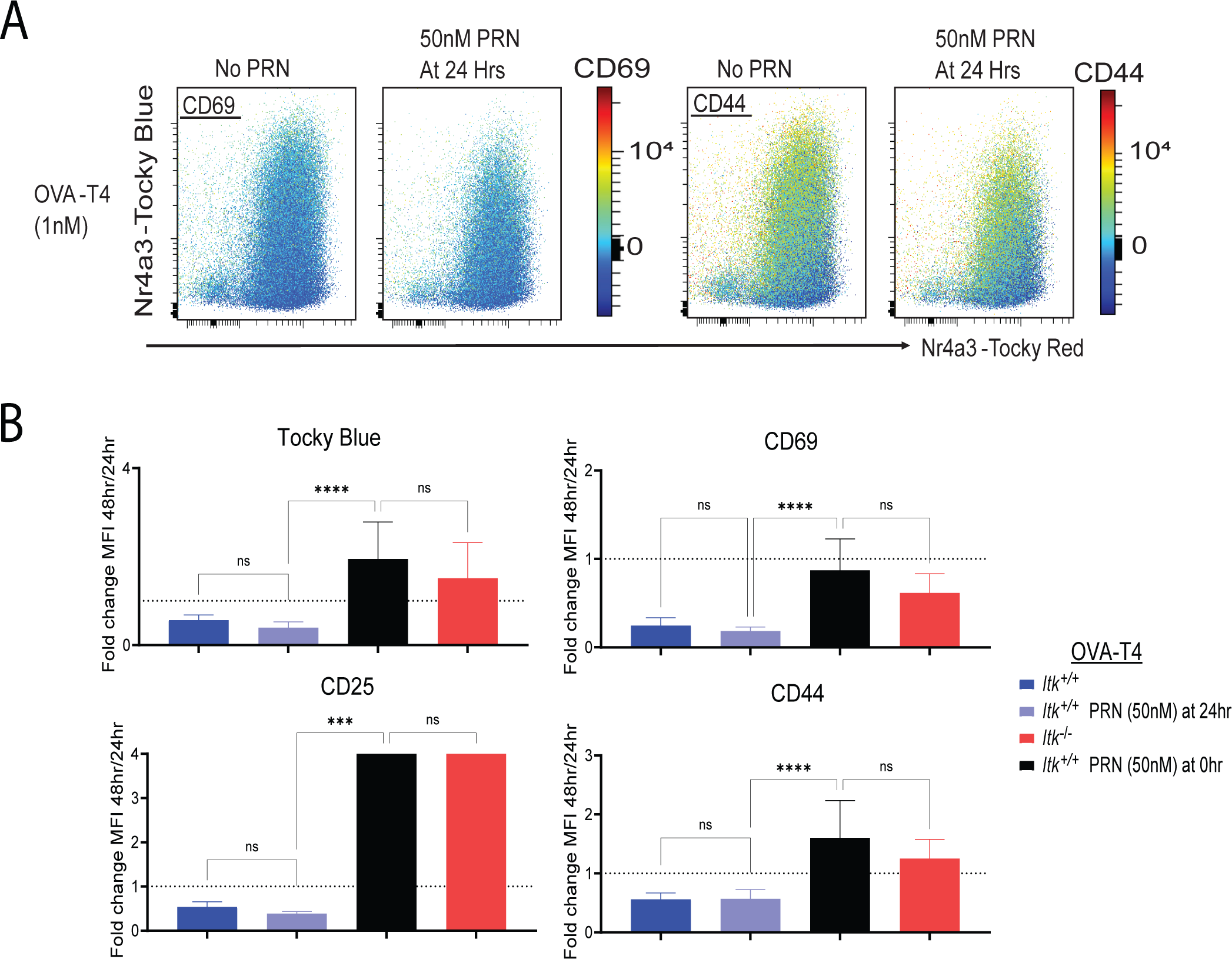
ITK signaling is required in the first 24h of activation to ensure signal termination at later times. (A) Tocky-Blue versus Tocky-Red fluorescence plots of OT-I *Nr4a3*-Tocky *Itk*^+/+^ cells stimulated for 48h with no PRN694, or with PRN694 (50nM) added at 24h. Colored continuous scale shows fluorescence of CD69 (left) or CD44 (right). (B) Compilation of fold change of normalized MFIs (% max of experiment) from 48h to 24h for CD69, Tocky-Blue, CD25, and CD44 stimulated with OVA-T4 (1nM). Statistics derived from one-way ANOVA with Dunn–Šidák correction, *p ≤ 0.05, **p ≤ 0.01, ***p ≤ 0.001, ****p ≤ 0.0001. Graph derived from 4-6 biological replicates from 4 experiments.

### Initial *Nr4a3-Tocky* expression is delayed and reduced in magnitude in the absence of ITK

TCR signal strength has been linked to the duration of TCR occupancy by pMHC ligands and is therefore a function of both the individual affinity of a TCR for pMHC as well as the density of pMHC ligands on the antigen-presenting cell (13–15, 31).

Together, these variables determine whether a T cell crosses the threshold to activation, and if activated, the time for induced transcriptional responses to begin (1, 26, 48, 49). One of the rate-limiting factors is the accumulation of second messengers, IP_3_ and DAG, generated through cleavage of PIP_2_ by PLC-γ1. For instance, a certain level of IP_3_ is needed to activate the calcium response, which leads to calcineurin activation, dephosphorylation of NFAT, and NFAT translocation into the nucleus, which takes place usually within minutes of TCR stimulation (32, 50, 51). Since ITK functions to promote PLC-γ1 activation, we reasoned that the absence of ITK should have a marked effect on the kinetics of upregulation of many T cell activation genes.

To test ITK’s role in the first 24h post stimulation, OT-I *Rag*1^-/-^ *Itk^-/-^ Nr4a3*-Tocky T cells were stimulated in parallel with OT-I *Rag*1^-/-^ *Itk^+/+^ Nr4a3*-Tocky T cells for 5h with OVA-N4 or OVA-T4 peptide. We first confirmed by bulk RNA-seq that the unstimulated starting populations of *Itk*^+/+^ and *Itk^-/-^* cells used for these experiments were not notably different in transcriptional profile due to potential effects of altered thymic development in the absence of ITK (Supp. Fig 3). To visualize the regulation of T cell surface receptors along with Tocky-Blue expression across multiple samples, we displayed the fluorescent signal of Tocky-Blue on the Y-axis and the MFI of the surface receptors (CD69 or CD25) in pseudo-color for each sample (Fig. 5A). This analysis showed that for *Itk*^+/+^ T cells stimulated with OVA-N4, Tocky-Blue expression was tightly coupled to both CD69 and CD25 upregulation. Stimulation with OVA-T4 produced a more heterogeneous response for both surface receptors as well as Tocky-Blue at 5h. *Itk^-/-^*T cells, in contrast, showed less synchronous responses, particularly after stimulation with OVA-T4. This was most apparent by assessment of CD25 upregulation, which was markedly reduced in Tocky-Blue-positive cells at 5h of stimulation (Fig. 5A). When comparing the expression of Tocky-Blue and CD69 in *Itk*^+/+^ and *Itk*^-/-^ cells, we observed the presence of CD69 positive cells that lacked Tocky-Blue expression in *Itk*^-/-^ cells (Fig 5B-C). After gating on CD69^+^ cells, only 50% of *Itk^-/-^* cells stimulated with OVA-T4 were positive for Tocky-Blue in comparison to the 80% of cells that were positive among CD69 expressing *Itk^+/+^* cells, further demonstrating a potential uncoupling of these two responses in the absence of ITK (Fig. 5C).

**Fig. 5.**
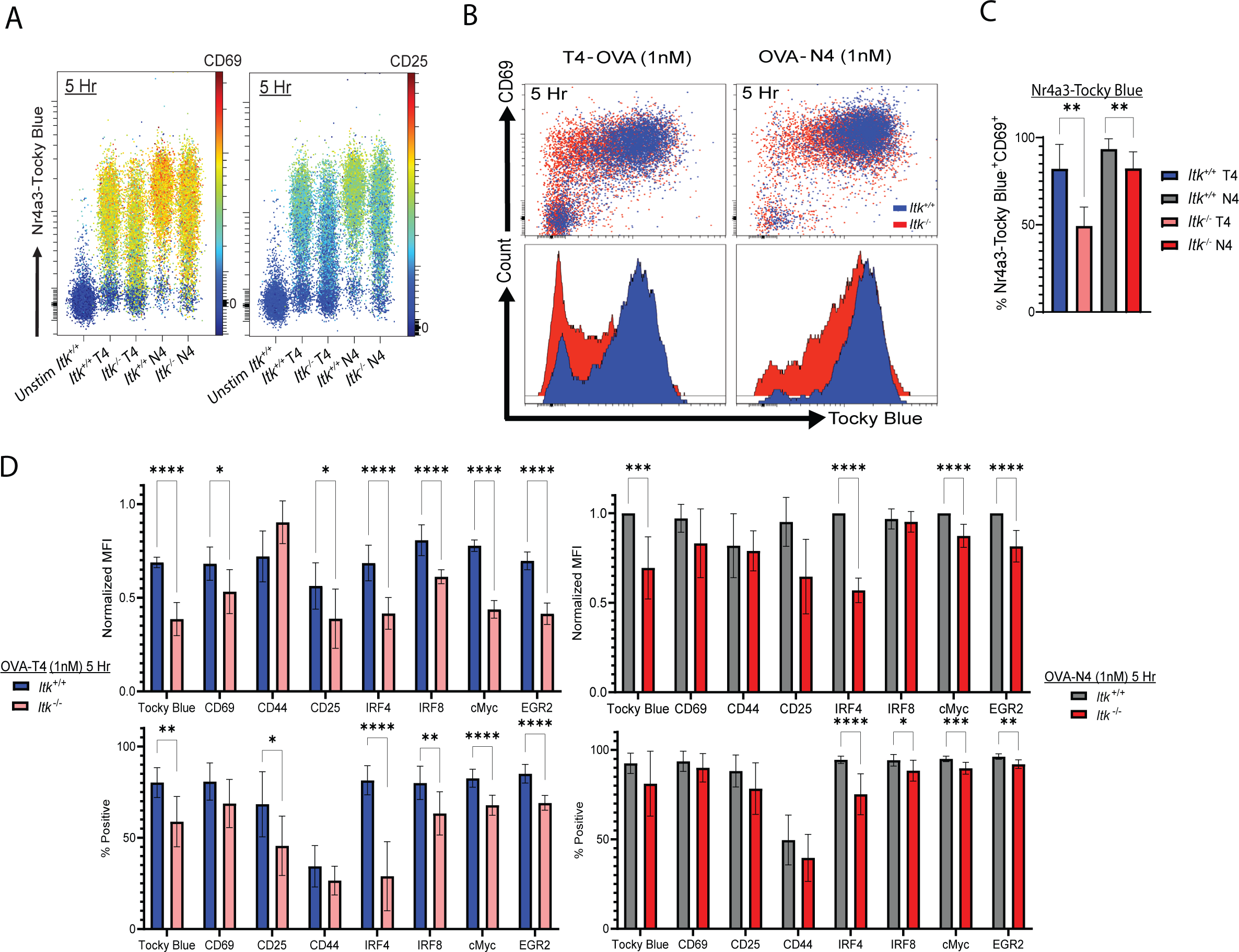
Signaling through ITK has varying contributions to the initial upregulation of distinct activation markers. (A) *Itk*^+/+^ versus *Itk*^-/-^ OT-I *Nr4a3*-Tocky cells were stimulated with OVA-N4 or OVA-T4 at 1nM for 5h. Comparative pseudocolor plots showing expression of Tocky-Blue fluorescence on Y-axis with CD69 or CD25 fluorescence on a colored continuous scale. (B) Pseudocolor plots of Tocky-Blue versus CD69 are shown with *Itk*^+/+^cells (Blue) overlaid on *Itk^-/-^* cells (Red) with histograms of Tocky-Blue fluorescence below. (C) Compilation of percentages of Tocky-Blue-positive cells of the CD69^+^ cells from 5 biological replicates and 4 experiments. Statistics derived from unpaired t-test, *p ≤ 0.05, **p ≤ 0.01, ***p ≤ 0.001, ****p ≤ 0.0001. (D) Compilation of data from 6-8 biological replicates of 6 experiments showing normalized MFI (as % experimental max MFI at 5h) (top) and percentages of positive cells (bottom) for a set of activation markers. Statistics derived from multiple unpaired t-test, *p ≤ 0.05, **p ≤ 0.01, ***p ≤ 0.001, ****p ≤ 0.0001.

Compiled data for a set of cell surface receptors and transcription factors shows the widespread effects of ITK deficiency on the early upregulation of TCR response genes (Fig. 5D). Transcriptional analysis of *Itk*^+/+^ versus *Itk^-/-^* cells stimulated with OVA-T4 for 5, 12, and 24h confirmed that the protein expression data correlated with mRNA levels for these TCR-induced genes (Fig. 3C, Supp. Fig 4). For most of these proteins, the percentages of cells with detectable upregulation and the expression levels per cell (i.e., MFIs) were reduced in the absence of ITK, an effect more clearly seen when cells were stimulated with the weaker peptide ligand OVA-T4 versus OVA-N4. At 5h post-stimulation, the expression levels per cell were far below the maximum expression achieved at later time points (Fig. 1D), indicating that the differences observed at this early time are primarily due to a delay in the onset of TCR-dependent transcriptional responses in the absence of ITK (Fig. 5D). Noting this differential expression early after TCR stimulation, we hypothesized that there were distinct subpopulations of cells that arose due to disparate thresholds of transcription factor activation being achieved by *Itk*^+/+^ compared to *Itk*^-/-^ T cells. To assess this, we performed dimensionality reduction by pooling flow cytometry data from *Itk*^+/+^ and *Itk*^-/-^ cells stimulated with OVA-N4 or OVA-T4 for 5h. Using the measurements of seven intracellular proteins and 3 extracellular proteins (Supp. Fig 5), we observed that *Itk^-/-^* T cells stimulated with 1nM T4 were highly enriched in cluster 1 over the other cell subsets. This cluster was characterized as CD69^+^ but low to negative for other induced proteins like IRF4, cMyc, IRF8, CD25, Egr2 and CD44, and likely represents cells that have achieved the threshold for upregulation of CD69, but not of the other proteins. Given the dependence of CD69 expression on Erk-MAPK signaling (43), these data highlight the marginal contribution of ITK to the rate of activation of the Erk-MAPK pathway compared to other transcriptional responses when CD8^+^ T cells are stimulated with low TCR signal strength (e.g., OVA-T4). These findings are consistent with our previous observations that ITK signaling is dispensable for early Erk-MAPK activation, but required for the normal rapid kinetics of NFAT activation (1). Here we expand our earlier findings to a larger panel of cell surface and intracellular markers, and show that ITK signaling impacts the early upregulation of many proteins, including *Nr4a3*-driven Tocky-Blue, CD25, CD44, cMyc, and Egr2, particularly under conditions of weaker TCR stimulation (Fig. 5D).

### Inhibition of NF**κ**B activation mimics the prolonged duration of TCR-dependent responses seen in the absence of ITK

Based on our findings that ITK was required during the first 24h of T cell stimulation to prevent the prolonged expression of multiple T cell activation markers, we considered that ITK might function to promote the upregulation of a gene (or genes) involved in the termination of TCR-dependent transcriptional responses. In our previous studies, we found that ITK had the most dramatic effect on NF-κB activation, as ITK is required for the optimal kinetics and magnitude of NF-κB(p65) nuclear translocation after TCR stimulation (1). In comparison, ITK signaling was necessary for the normal rapid kinetics of NFAT translocation into the nucleus but not for the magnitude of NFAT activation, and had little effect on the kinetics of Erk-MAPK phosphorylation (1). To address the possibility that one or more of these transcriptional pathways might contribute to the altered kinetics of gene expression observed in *Itk^-/-^* T cells, we performed the same time course experiments in the presence or absence of three different inhibitors: an inhibitor of NFAT activation, Cyclosporin A (CsA); an inhibitor of NFkB activation, IKK16; or an inhibitor of Erk-MAPK phosphorylation, PD325901 (a MEK inhibitor). The outcome seen for each of these inhibitors was compared to the effects of germline KO of ITK with particular attention being paid to the later time points where *Itk*^+/+^ cells show termination of signaling and loss of activation marker expression compared to *Itk^-/-^*T cells. These experiments showed that *Itk^+/+^* T cells treated with IKK16 experienced the prolonged TCR signaling that is seen in T cells lacking ITK, evident by the continued co-expression of Tocky-Blue and CD69 at 48h with OVA-N4 stimulation compared to *Itk*^+/+^(Fig. 6A). We also observed a similar prolonged expression of several other proteins when comparing IKK16-treated T cells to *Itk^-/-^* T cells, including CD44, CD69, Tocky Blue, Egr2, cMyc and IRF4 (Fig. 6B-D). At 5h post stimulation IKK16 inhibited CD8+ T cells showed a robust decrease from *Itk*^+/+^ and *Itk*^-/-^ in expression of most early activation markers with both high and low affinity pMHC stimulation (Fig. 6E). At 5h IKK16 inhibited and *Itk*^-/-^ CD8 T cells showed similar levels of expression for many markers including Tocky Blue, CD25, IRF4, cMyc and Egr2 with both OVA-T4 and OVA-N4 stimulation (Fig. 6E). These data strongly suggest that ITK signaling is required for the upregulation of an NF-κB-dependent response gene that terminates/tunes TCR induced gene expression in *Itk^+/+^*CD8^+^ T cells. We also observed a higher peak magnitude of expression of the NFAT-dependent *Nr4a3*-Tocky Blue protein in the IKK-16 treated *Itk*^+/+^ cells compared to untreated at 24h (Fig. 6B). This finding further supports the existence of an NF-κB-dependent mechanism that tunes the NFAT response in the timeframe of 12-24h post-stimulation.

**Fig. 6.**
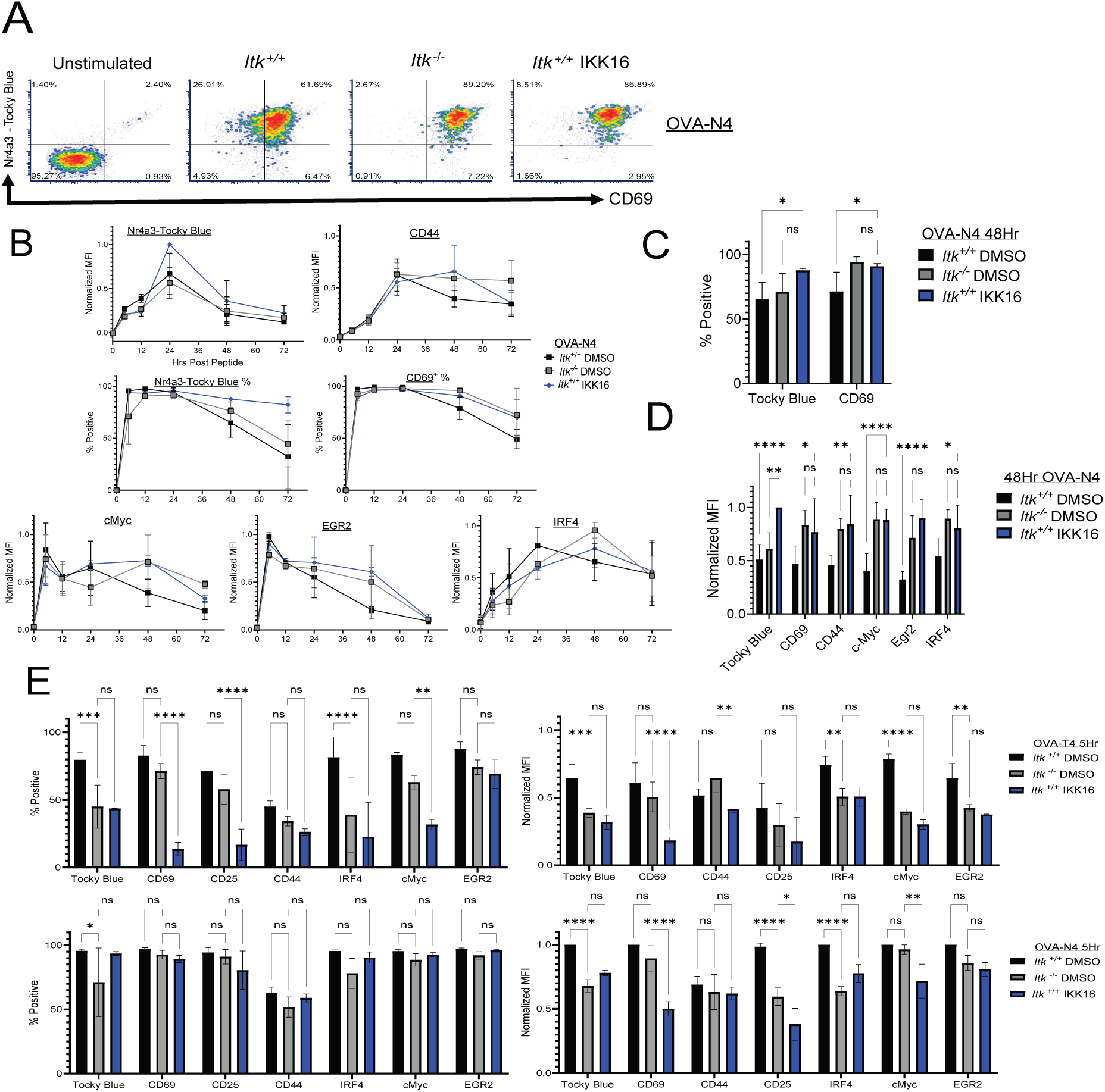
Inhibition of NF-κB activation leads to prolonged expression of *Nr4a3*-Tocky-Blue. (A) Tocky-Blue versus CD69 fluorescence at 48h in OT-I *Nr4a3*-Tocky *Itk*^+/+^ cells, *Itk^-/-^* cells, and *Itk*^+/+^ cells stimulated with 1nM of OVA-N4. *Itk*^+/+^ cells were stimulated in the presence or absence of inhibitors of NF-κB activation (IKK-16 (B) Compiled data showing expression (MFI and/or % positive, as indicated) of Tocky-Blue, CD69, PD-1, IRF4, cMyc, and Egr2 for time courses of 0-72h for cells cultured in the presence or absence of IKK16; all cells were stimulated with 1nM OVA-N4. (C) Statistics derived for percentages of marker positive CD8^+^ T cells at 48h post Ova addition comparing *Itk*^+/+^, *Itk^-/-^*, and *Itk*^+/+^ cells treated with IKK16. Statistics derived from two-way ANOVA with Tukey’s correction, *p ≤ 0.05, **p ≤ 0.01, ***p ≤ 0.001, ****p ≤ 0.0001 using 3 biological replicates from 2 experiments. (D) Statistics of normalized MFI (% of 48h max) in CD8^+^ T cells at 48h post Ova addition comparing *Itk*^+/+^, Itk*^-/-^* , and *Itk*^+/+^ inhibited with IKK16. Statistics derived from two-way ANOVA with Tukey’s correction *p ≤ 0.05, **p ≤ 0.01, ***p ≤ 0.001, ****p ≤ 0.0001 using 3 biological replicates from 2 experiments. (E) Graphs show compilations of normalized MFI (% experimental max of marker at 5h (Right) and percentages of positive cells (Left) for *Itk^-/-^* cells and *Itk*^+/+^ treated with IkK inhibitor (IKK16) for 5h with OVA-T4 (Top) or OVA-N4 (Bottom). Data are mean +/-SD of 3 biological replicates over 2 experiments. Statistics derived from two-way ANOVA with Tukey’s correction, *p ≤ 0.05, **p ≤ 0.01, ***p ≤ 0.001, ****p ≤ 0.0001 using 3 biological replicates from 2 experiments. (F) Mathematical modeling of IRF4 expression over time (hrs) showing expression and promoter activity when stimulated with high affinity OVA-N4.

### Inhibition of NFAT leads to increased rate of expression of MAPK-ErK dependent markers

In contrast to the effects of IKK-16, inhibition of NFAT activation with CsA or of Erk-MAPK phosphorylation with PD325901 had varying effects on the kinetics and/or the peak expression levels of multiple activation markers; however, neither of these inhibitors led to prolonged TCR signaling or sustained expression of several proteins (Fig. 7A). Instead, NFAT inhibition with CsA often mimicked the reduced upregulation of proteins such as CD69, CD44, CD25, IRF4, and cMyc at 5h post-stimulation as seen in the *Itk^-/-^* (Fig. 7B). To examine early kinetics, we used expression levels of Nr4a3-Tocky Blue as pseudo-time since initial TCR signaling, gating on increasing levels of the Nr4a3-Tocky Blue reporter. This analysis showed that Erk-MAPK-dependent CD44, CD25, and CD69 upregulation all increased at a faster rate in *Itk*^+/+^ cells treated with Cyclosporin A compared to untreated cells (Fig. 7C). ITK inhibition has been shown to delay translocation of NFAT into the nucleus through regulation of signaling thresholds proximal to TCR signaling (1) which could explain the overall increase in CD44 at 5Hrs between *Itk*^-/-^ and *Itk*^+/+^ at 5h when stimulated with low affinity OVA-T4 (Fig. 5D).

**Fig. 7.**
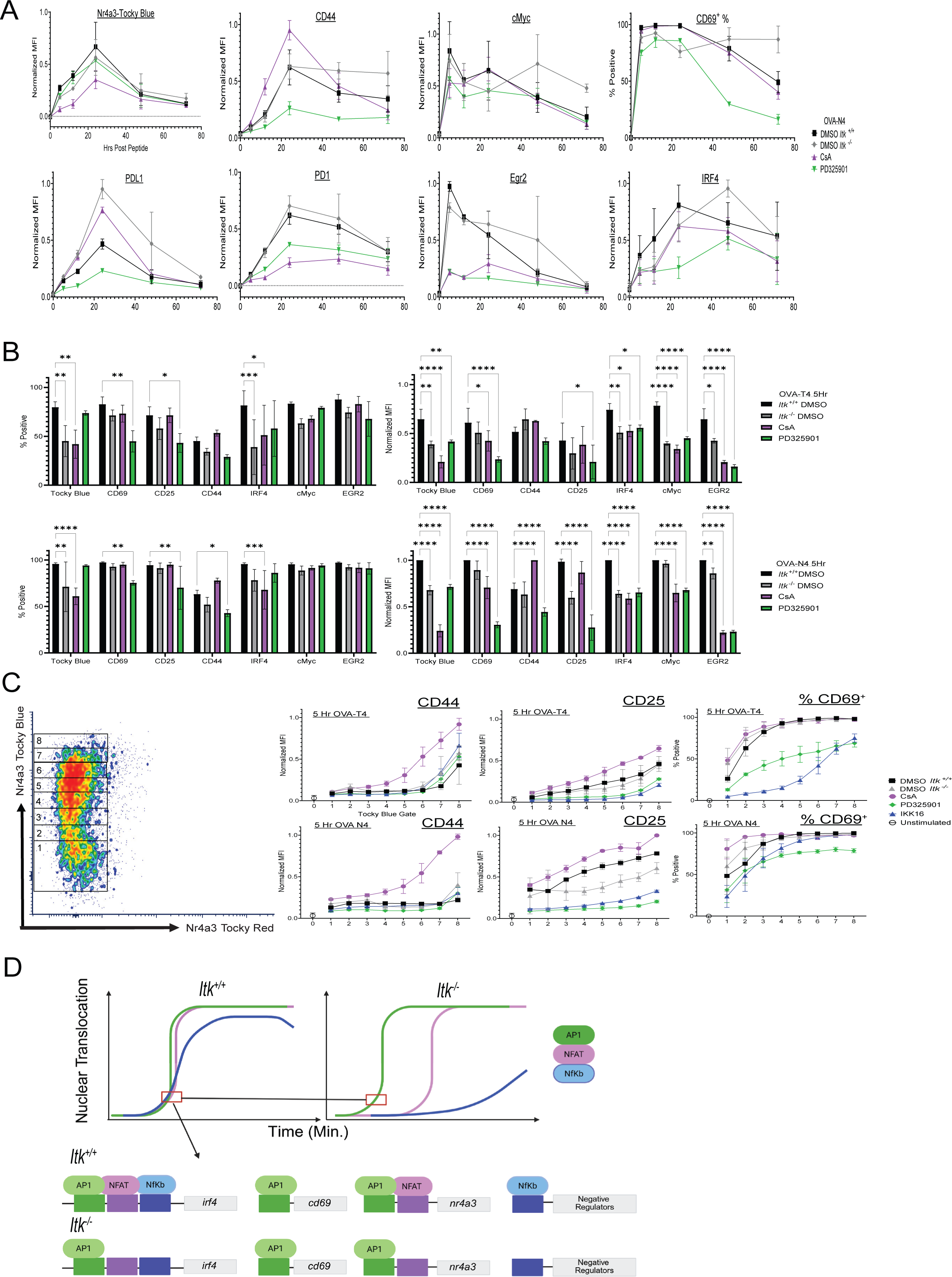
Inhibition of NFAT activation leads to increased expression of MAPK-ErK dependent activation markers. (A) Compiled data show expression (MFI and/or % positive, as indicated) of Tocky-Blue, CD44, CD69, PDL1, PD-1, Egr2 and IRF4 for time courses of 0-72h. *Itk^-/-^* cells and *Itk*^+/+^ cells cultured in the presence or absence of CsA or PD325901 are shown; all cells were stimulated with 1nM OVA-N4. Data are mean +/-SD; representative of 3 biological replicates over 2 experiments. (B) Graphs show compilations of normalized MFI (% experimental max of marker at 5h (Right) and percentages of positive cells (Left) for *Itk^-/-^* cells and *Itk*^+/+^ treated with calcineurin inhibitor (CsA) or MEK Inhibitor (PD325901) for 5h with OVA-T4 (Top) or OVA-N4 (Bottom). Data are mean +/-SD of 3 biological replicates over 2 experiments. (C) Pseudo-time kinetics shown by normalized MFI (ratio of marker MFI to experimental max at 5h) of CD44 and CD25, and % cells positive for CD69, after gating on different levels of Tocky Blue. Representative gating scheme for different levels of Tocky Blue are shown with the density plot at left. Data are compiled from 3 biological replicates from 2 experiments. (D) Model illustrates the proposed differential contribution of transcription factors NFAT, NFκB, and AP-1 to genes encoding IRF4, CD69, and Nr4a3, and the impact of ITK signaling on these pathways, including the induced expression of a putative negative regulator of TCR induced gene expression. Made with Biorender.

Moreover, this finding suggests that in the absence of NFAT activation, genes that are predominantly dependent on the Erk-MAPK pathway are more rapidly induced compared to cells where both NFAT and the MAPK-Erk pathway are rapidly induced showing another avenue at which temporally dysregulated processes can lead to differential gene regulation (Fig. 7D).

## Discussion

Our findings provide new insights into the function of the Tec family kinase, ITK, in regulating TCR-dependent gene expression. Using a dynamic fluorescent reporter to capture information about TCR signaling kinetics along with the assessment of multiple cell surface receptors and transcription factors, we show that ITK signaling contributes to both early and late T cell activation responses. At early time points after TCR stimulation, ITK is required for the rapid upregulation and peak expression levels of many TCR-induced proteins. In addition, our data suggest that in the first 24 hoursof activation, ITK induces a regulatory pathway that is needed at later times to terminate the expression of many *trans* regulatory pathway dependent markers such as CD69, CD44, and IRF4.

These findings indicate that the characterization of ITK as a kinase that regulates ‘TCR signal strength’ is an oversimplification of ITK’s role in T cell activation. At early timepoints post stimulation, CD8^+^ T cells lacking ITK often show delayed and impaired responses that resemble those of *Itk*^+/+^ cells responding to low affinity TCR:pMHC interactions. We observed this in our earlier studies using bulk RNA-seq to compare gene expression in *Itk*^+/+^ OT-I cells stimulated *in vitro* with OVA-N4 or OVA-T4 in the presence or absence of the dual ITK/RLK inhibitor PRN694 (1). Examining very early transcriptional responses at 0.5, 1, and 2h post-stimulation we saw multiple examples of genes (e.g., *fos, il2, il2ra, irf4, ifng, nfkbid, nfkbiz*) that fit this pattern. However, this pattern of response did not apply to the majority of proteins examined in our study (Fig. 3B, Supplemental Fig 1). Our prior studies demonstrated a differential role for ITK in the activation of NFAT, NFκB (p65), and Erk-MAPK pathways (1). Based on our inhibitor data shown here, we found that each gene examined showed differential dependence on the three major PLCγ1-dependent pathways (Fig. 6B-E, 7A-C), explaining the varying impact of ITK signaling on the panel of proteins we analyzed. Given the ongoing efforts to develop therapeutic ITK inhibitors for autoimmune diseases and immunotherapies, our results indicate an ongoing need to dissect the detailed consequences of blocking ITK signaling.

The duration of TCR signaling is regulated through many different mechanisms. These include ubiquitination and degradation of key signaling proteins (52–54), the recruitment of protein tyrosine and lipid phosphatases (55, 56) to the immune synapse, and enzymes that inactivate the PLCγ1-dependent second messengers IP_3_ and DAG (16, 57, 58). More complex mechanisms, such as negative feedback loops that are mediated by post-translational modifications of signaling intermediates have also been described (59). In addition to these relatively rapid proximal signaling events, other inhibitory factors including Suppressor of Cytokine Signaling (SOCS) family proteins are absent in naïve cells and rapidly increase in expression upon TCR activation (60). Less well-studied are the negative regulators of TCR signaling that are transcriptionally induced following T cell stimulation. While the specific details of the ITK-dependent regulatory mechanism identified here remain to be elucidated, our results strongly suggest that an NFκB-regulated gene is contributing to the prolonged signaling phenotype. ITK has been shown to play a substantial role in NFκB activation compared to Erk-MAPK-dependent and NFAT-family transcription factors (1). This is likely due to ITK’s importance in activating PLC-γ1 and the known roles for pathways downstream of both DAG and IP_3_ on the nuclear translocation of NFκB. We hypothesize that the absence of ITK leads to reduced NFκB activation and thus to the impaired expression of one or more negative regulators of TCR signaling. This hypothesis is supported by the elevated expression of NFAT-dependent *Nr4a3*-Tocky Blue at 24h and prolonged expression of CD44, CD69, cMyc and Egr2 at 48hs in cells treated with IKK-16, effects not observed in cells treated with Cyclosporin A or PD325901.

Our detailed studies of the kinetics of T cell activation-dependent gene expression responses reveal the importance of clearly defining ‘T cell activation’ in experimental settings. For instance, CD69 has historically been used as an activation marker due to its rapid expression after T cell stimulation. In our time-course experiments, we showed that both CD69 and Nr4a3-Tocky-Blue were coordinately co-expressed in both *Itk*^+/+^ and *Itk*^-/-^ CD8^+^ T cells following strong TCR stimulation. However, lower affinity stimulation uncoupled these two responses, with a significant proportion of *Itk^-/-^*OT-I cells upregulating CD69 without expressing Tocky-Blue at 5h. As *Cd69* is known to be an Erk-MAPK-dependent gene, this finding is consistent with observations showing that the Erk-MAPK pathway has a lower threshold for activation than that needed for NFAT activation and is relatively unaffected by the absence of ITK activity (1, 3, 5, 61).

Unsupervised clustering analysis performed using activation marker expression further corroborated these findings, as the unique clusters 1, 3, and 4 were dominated by OVA-T4 stimulated *Itk^-/-^* cells and was enriched for several Erk-MAPK target genes, such as Egr2, CD69, cMyc and IRF8 (Supp. Fig 5) (60, 62, 63). Overall, these findings strongly suggest that T cells stimulated in the absence of ITK have a prolonged lag time between the initiation of the Erk-MAPK pathway and the activation of NFAT (Fig. 7A-D) A recent study used dual-reporter-positive T cells for simultaneous assessment of NFAT and Erk-MAPK nuclear translocation. The authors showed that over long timeframes (∼30h), these two transcription factors oscillate between the nucleus and the cytoplasm in a peptide affinity- and dose-dependent manner (37) given continuous pMHC-TCR interaction, providing a mechanism to generate divergent transcriptional programs in cells stimulated under different conditions. Furthermore, they showed through mathematical modeling and plate bound pMHC stimulation that when AP-1 and NFAT have DNA binding availability at the same time they act in a way that keeps them from binding alone or to other binding partners. Based on our data, we anticipate that the absence or inhibition of ITK would alter this oscillatory behavior and significantly impact the T cell programming observed in this study, particularly if the sequence at which AP-1 and NFAT translocation into the nucleus is delayed through ITK inhibition or low pMHC binding affinity. One observation of interest was our finding that inhibiting NFAT activation through Cyclosporin A treatment led to faster and increased expression of CD44, CD25, and CD69 compared to control cells (Fig. 7 F-H). Based on the robust inhibition of CD69, CD44, and CD25 expression by the MEK inhibitor PD325901, it is apparent that these proteins are highly dependent on Erk-MAPK and likely AP-1 activities. One possibility is that inhibition of NFAT nuclear translocation results in elevated AP-1 activity due to the absence of the NFAT/AP-1 complexes competing for nuclear AP-1. As a result, genes that are regulated by AP-1, but not NFAT/AP-1, would be more readily occupied by higher levels of AP-1. Together, these findings demonstrate the important insights to be gained by detailed molecular and biochemical analyses of early responses to TCR signaling. Unraveling these pathways reveals the complex interplay between transcriptional regulators acting to induce downstream effector genes, their abilities to act solo or in tandem with others, and their essential roles in regulating signal termination.

### Experimental Methods Mice

Mice were housed and bred at the University of Colorado-Anschutz campus in accordance with IACUC (Institutional Animal Care and Use of Committee guidelines.) Previously described *Itk^-/-^*(64) mice were bred to C57BL/6-Tg(TcraTcrb)1100Mjb/J - *Rag*1tm1Mom/J (Jackson Laboratory) to generate OT-I *Rag1*^-/-^ *Itk*^-/-^. Wild-type or *Itk*^-/-^ OT-I *Rag1*^-/-^ mice were then crossed to the *Nr4a3*-Tocky Timer line (38). The *Nr4a3*-Tocky reporter was bred to homozygosity, and mice were genotyped using a SYBR green protocol (42).

### OT-I *Nr4a3*-Tocky CD8^+^ T cell Stimulation

Spleens were harvested and pooled from 6-12 week old OT-I *Rag*1^-/-^ *Nr4a3*-Tocky and OT-I *Rag1*^-/-^ *Nr4a3*-Tocky *Itk^-/-^* mice. In some experiments, splenocytes were stained with 0.14ng/uL CFSE (Life Technologies) for 20 minutes in dPBS then washed twice with flow buffer (dPBS + 2% FBS) prior to stimulation. 2 x 10^6^ splenocytes per well were stimulated with SIITFEKL or SIINFEKL peptides (21^st^ Century Biochemicals) in the presence of 2μg/mL LPS (Sigma Aldrich) and incubated at 37C. Cells were harvested for spectral flow cytometry analysis at 5, 12, 24, 48, and 72 hours post-peptide addition. For longer timepoints (48, 72h), peptide and LPS were washed off at 24h and splenocytes transferred into new plates and given fresh media supplemented with 10 IU mIL-2 (Peprotech). Media supplemented with 10 IU mIL-2 was replaced again at 48 hours. In some experiments the small molecule inhibitor of ITK/RLK, PRN694 (Sigma), was added at 50nM at time 0 or at 24h post Ova + LPS addition following washing. In other experiments, Cyclosporin A (Sigma) at 200ng/mL, IKK-16 (Sigma) at 500nM, or PD325901 MEK inhibitor (Tocris Bioscience) at 1μM were added to splenocytes 20 minutes before the Ova peptide + LPS addition.

### Flow Cytometry Analysis

Bulk splenoyctes were harvested and stained with a mixture of Anti-mouse CD16/CD32 Fc Shield (Clone 2.4G2) at 0.5ug/mL and Ghost Dye Red 780 Viability Dye (diluted 1:15000) for 15 minutes at 4C before washing with PBS containing 2% FBS (Cytiva).

Surface marker master mix was made with Brilliant Stain Buffer (Cytiva) diluted 1:2 in dPBS (Gibco) and cells were stained in 100uL for 20 minutes at 4C. For intracellular panels, cells were further washed then fixed and permeabilized prior to antibody staining. Antibodies used were as follows: CD8-BUV395 (Clone 53-6.7), CD69-BUV805 (Clone: H1.2F3), CD44-AF700 (Clone: IM7), PD-1-FITC (Clone: 29F.1A12), CD25-PE (Clone: PC61.5), , CD122-PE-Cy7 (Clone: TM-B1), CD274 (PDL1)-BUV737 (Clone:MIH5), CD127 (Clone: A7R34), CD62L-Percp-Cy5.5 (Clone: MEL-14), IRF4-AF594 (Clone: IRF4.3E4), c-Myc-AF647 (Clone: E5Q6W), Egr2 (Clone: erongr2), Eomes-BUV737 (X4-83), IRF8 (Clone: V3GYWCH), TCF1/7-BV421 (Clone :S33-966), IkB-AF488 (Clone:L35A5).

### Bulk mRNA-Sequencing Sample Preparation

Cells from each genotype (OT-I *Rag1*^-/-^ *Itk*^+/+^ *Nr4a3*-Tocky^+/+^ and OT-I *Rag1*^-/-^ *Itk*^-/-^ *Nr4a3*-Tocky^+/+^) were separately pooled from spleen and lymph nodes (inguinal, axillary) and red blood cells were lysed with ACK lysis buffer. For unstimulated samples, naive T cells were enriched with the StemCell EasySep Mouse CD8^+^ T cell Isolation Kit according to manufacturer’s instructions and sorted for live, CD44^lo^ CD8^+^ naïve cells..

For all other time points (5, 12, and 24 hours), isolated cells were split into two groups. With one group, naïve CD8+ T cells were isolated with the Stemcell EasySep Mouse Naïve CD8+ T cell Isolation Kit, according to manufacturer’s instructions. From the rest of the cells, T cells were depleted using the Stemcell EasySep Mouse CD90.2 Positive Selection Kit II, according to manufacturer’s instructions. Naïve T cells were plated with T-depleted cells (treated as APCs) in T cell media at a ratio of 5:1 in a 24-well plate with either 1nM OVA-N4 or OVA-T4 peptide. After 5, 12, and 24 hours of culture, cells were harvested and stained with Zombie Green and CD8-PE (clone 53-6.7). Cells were sorted on a Bio-Rad S3e Cell Sorter, gated on single cells, live, CD8+ T cells. Sort efficiency was verified via flow cytometry. Sorted CD8+ T cells were washed once in cold PBS before being resuspended in 350ul of buffer RLT +3.5ul BME and stored at - 80 until RNA isolation. After all samples had been acquired, frozen CD8+ T cell lysates were thawed and total RNA was isolated using the RNeasy Mini Kit (Qiagen) according to manufacturer’s instructions. On-column DNA digestion was performed using the Qiagen RNAse-free DNase set, and RNA cleanup was performed before RNA quantification on a NanoDrop. RNA was frozen at -80 and sent to National Jewish Health Genomics Core for library preparation and sequencing.

### Watchmaker mRNA Library Prep and Sequencing

RNA sequencing libraries were prepared according to the Watchmaker mRNA Library Build user guide. Briefly, mRNA from 5ng of total RNA was isolated using polyA, oligo-dT magnetic beads. The isolated mRNA was then subject to enzymatic fragmentation, resulting in ∼200 bp fragments. The resulting RNA fragments underwent first and second strand cDNA synthesis. Unique KAPA Dual-Indexes were then ligated to the cDNA. The ligated product was then PCR amplified for 16 cycles. The resulting libraries were quantified using the Qubit HSDNA assay and the TapeStation HSDNA 1000 assay. Equal molar concentrations were pooled, diluted, and sequenced using a 200 cycle Illumina NextSeq 2000 flow cell.

### Processing and Analysis of Bulk mRNA-Sequencing Reads

Adapter sequences were trimmed from quality raw sequences reads with cutadapt version 4.2 (65) and then aligned to mm39 mouse reference genome with STAR version 2.7.10b (66). Samples were filtered to retain expressed genes (at least 20 reads present in at least half of the samples). Differential expression analysis was performed with DESeq2 version 1.45.3 (67) to identify induced genes (stimulated conditions versus unstimulated controls). Processing of the sequencing reads was done using the Alpine high performance computing resource at the University of Colorado Boulder Alpine Research Computing (68). Analysis performed within R version 4.4.1, data visualized using ggplot2 (69).

### Dimensionality Reduction

Unsupervised clustering was done in Flowjo (version 10.10). Gates were set for Live CD8+ T cells and downsample (version 3.3.1) was ran to 3000 cells per sample.

Samples were concatenated and tSNE was ran through the flowjo native platform for activation markers. Xshift (version 1.4.1) (70) was used for automated clustering for all activation markers and clusters were visualized within Flowjo native platform cluster explorer.

### Mathematical Modeling

To model the dynamics of the Tocky-Blue signal post peptide stimulation we use the following ordinary differential equation model

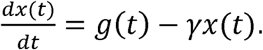

Here *x*(*t*) is the signal at time *t* post stimulation, and given the reported ∼4.1h half-life of the Tocky-Blue protein (38), the degradation rate is set as 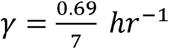. The function *g*(*t*) represent the *Nr4a3* promoter activity that is assumed to take the form

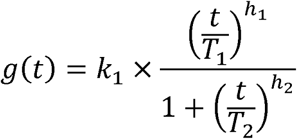

with positive constant parameters *k*_1_, *r*_1_, *h*_1_, *r*_2_, *h*_2_. The specific values of these constants determine the shape of *g*(*t*). If the exponent *h*_1_ is smaller than *h*_2_, then this function captures promoter activity that starts at zero and first increases with time *t* to reach a maximum, and then decreases back to zero with increasing *t*. For each peptide ligand and cell genotype, the five parameters defining *g*(*t*) were inferred by performing a least-square error optimization between the experimentally observed average normalized MFI for Tocky Blue at 5h, 12h, 24h, 48h and 72h post-stimulation, and the corresponding model predictions at these time points.

## Supporting information

Supplement Figures Combined

## Acknowledgements

We thank the Barbara Davis Center BioResource/Molecular Core Center at the University of Colorado-Anschutz for flow cytometry support, and the office of Laboratory Animal Resources excellent mouse work. We also thank Loni Perrenoud for expert assistance in flow cytometry data analysis. **Funding:** This work was supported by grants from the NIH (AI120423 and R01AI043957), and the US-Israel Binational Science Foundation (2021035). AS acknowledges the support of NIH-NIGMS via grant R35GM148351. **Author Contributions:** J.H planned optimized and performed the experiments, analyzed flow cytometry data, and wrote the manuscript. Z.B. performed the bulk mRNA sequencing experiments and analyzed the bulk mRNA seq data. U.C. helped with analysis, revisions, and editing. A.R. performed experiments, revisions and editing. M.M contributed to manuscript revisions and editing. M.O created the Nr4a3-Tocky mice. A.S created all of the kinetic mathematical models and edited the manuscript. L.B. directed all experiments, assisted with data analysis and interpretations, and edited the revised the manuscript. **Competing Interests, Data and Material Availability:** the authors declare no competing interests relating to this work. All data generated and material used are available upon request.

---

**Supplemental Fig. 1. Time course of expression of T cell activation markers in *Itk*^+/+^ versus *Itk^-/-^* T cells reveals distinct patterns of ITK-dependence.**

Compilation of data from 0-72h timecourses show normalized MFI for each protein in cells expressing that protein. Data are mean +/-SD of normalized MFI (% experimental max) from 3-8 biological replicates in 3-5 experiments.

**Supplemental Fig. 2 Mathematical modeling of protein expression estimates prolonged promoter activation of Nr4a3-Tocky Blue in *Itk*^-/-^ CD8^+^ T cells**

Mathematical modeling showing estimated expression and promoter activation of *Nr4a3* in *Itk*^+/+^ and *Itk*^-/-^ cells when stimulated with low affinity OVA-T4 and high affinity Ova-N4 over the time course of 0-72h.

**Supplemental Fig. 3. Principle component analysis verifies that *Itk*^+/+^ and *Itk^-/-^* OT-I cells are transcriptionally similar upon isolation.**

*Itk*^+/+^ and *Itk^-/-^* OT-I cells were stimulated for 5, 12, and 24h with 10nM OVA-T4 then sorted for CD8^+^ T cells prior to bulk RNA-seq analysis. Unstimulated cells were analyzed in parallel as a control. Principle component analysis on 2-3 independent samples was performed.

**Supplemental Fig. 4. Bulk RNA-seq analysis shows reduced transcription of T cell activation-induced genes in the absence of ITK.**

*Itk*^+/+^ and *Itk^-/-^*OT-I cells used for all studies were stimulated for 0, 5, 12, and 24h with OVA-T4 at 10nM, and then sorted for CD8^+^ T cells prior to bulk RNA-seq analysis. Graphs show transcript numbers (per million) for the indicated genes.

**Supplemental Fig. 5. Unsupervised cluster analysis of intracellular markers shows delayed upregulation of genes in *Itk^-/-^*T cells.**

Unsupervised clustering was performed on pooled cells from *Itk*^+/+^ and *Itk^-/-^* OT-I *Nr4a3*-Tocky T cells stimulated with 1nM OVA-N4 or 1nM OVA-T4 for 5h. Samples were downsampled to 3000 CD8^+^ T cells per condition prior to concatenation. TSNE pseudocolor plot shows distribution of clusters (left) and bar graph (right) show the percentages of cells in each cluster. Histograms (middle row) show normalized cell numbers present in each cluster. Heat map (bottom) shows relative expression of each protein within each cluster.

