## Supplement Figures Combined for "Regulation of TCR-induced Transcriptional Kinetics by Interleukin 2-inducible T Cell Kinase (ITK) in CD8^+^ T cells"

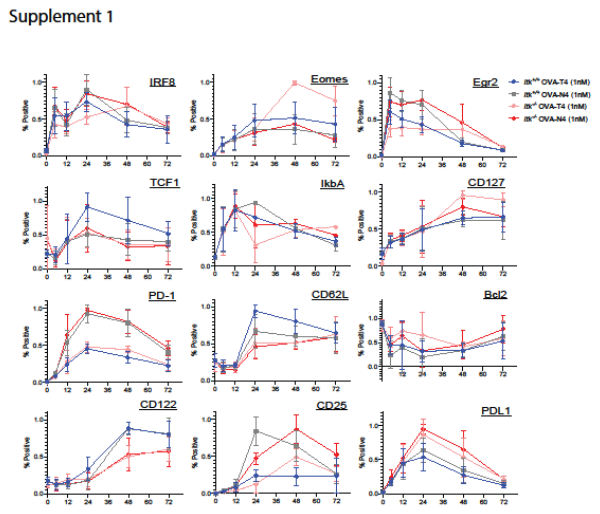

**Supplemental Fig. 1. Time course of expression of T cell activation markers in *Itk*^+/+^ versus *Itk^-/-^* T cells reveals distinct patterns of ITK-dependence.**

Compilation of data from 0-72h timecourses show normalized MFI for each protein in cells expressing that protein. Data are mean +/-SD of normalized MFI (% experimental max) from 3-8 biological replicates in 3-5 experiments.

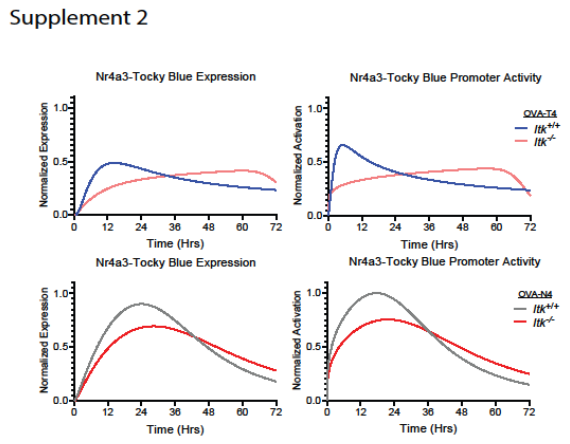

**Supplemental Fig. 2 Mathematical modeling of protein expression estimates prolonged promoter activation of Nr4a3-Tocky Blue in *Itk*^-/-^ CD8^+^ T cells.**

Mathematical modeling showing estimated expression and promoter activation of *Nr4a3* in *Itk*^+/+^ and *Itk*^-/-^ cells when stimulated with low affinity OVA-T4 and high affinity Ova-N4 over the time course of 0-72h.

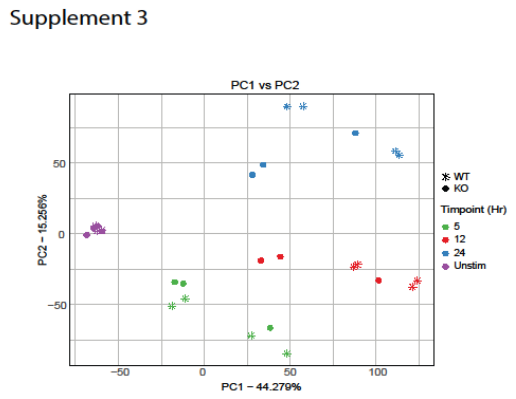

**Supplemental Fig. 3. Principle component analysis verifies that *Itk*^+/+^ and *Itk^-/-^* OT-I cells are transcriptionally similar upon isolation.**

*Itk*^+/+^ and *Itk^-/-^* OT-I cells were stimulated for 5, 12, and 24h with 10nM OVA-T4 then sorted for CD8^+^ T cells prior to bulk RNA-seq analysis. Unstimulated cells were analyzed in parallel as a control. Principle component analysis on 2-3 independent samples was performed.

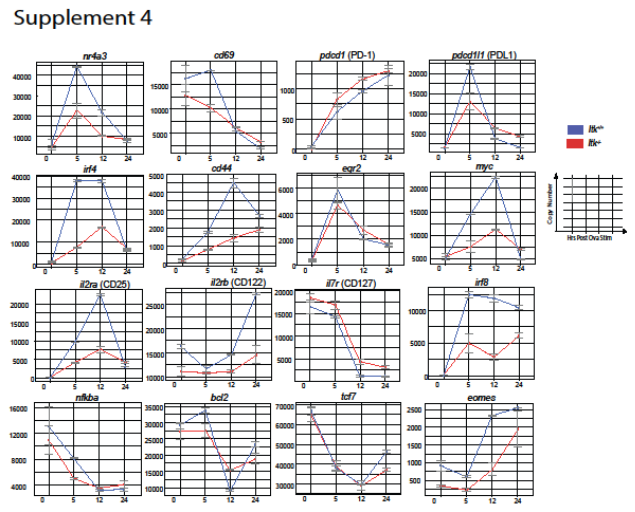

**Supplemental Fig. 4. Bulk RNA-seq analysis shows reduced transcription of T cell activation-induced genes in the absence of ITK.**

*Itk*^+/+^ and *Itk^-/-^* OT-I cells used for all studies were stimulated for 0, 5, 12, and 24h with OVA-T4 at 10nM, and then sorted for CD8^+^ T cells prior to bulk RNA-seq analysis. Graphs show transcript numbers (per million) for the indicated genes.

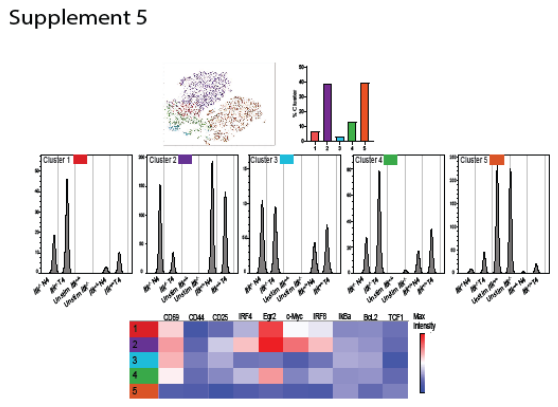

**Supplemental Fig. 5. Unsupervised cluster analysis of intracellular markers shows delayed upregulation of genes in *Itk^-/-^* T cells.**

Unsupervised clustering was performed on pooled cells from *Itk*^+/+^ and *Itk^-/-^* OT-I *Nr4a3*-Tocky T cells stimulated with 1nM OVA-N4 or 1nM OVA-T4 for 5h. Samples were downsampled to 3000 CD8^+^ T cells per condition prior to concatenation. TSNE pseudocolor plot shows distribution of clusters (left) and bar graph (right) show the percentages of cells in each cluster. Histograms (middle row) show normalized cell numbers present in each cluster. Heat map (bottom) shows relative expression of each protein within each cluster.
